# Culture filtrate selectively promotes *Mycobacterium tuberculosis* growth from extremely low density inocula: implications for quantifying differentially culturable phenotypes

**DOI:** 10.1101/2025.10.15.682508

**Authors:** Ryan Dinkele, Ditshego Ralefeta, Atica Moosa, Digby F. Warner, Sophia Gessner

## Abstract

Differentially culturable (DC) *Mycobacterium tuberculosis* (*Mtb*) phenotypes reduce the sensitivity of sputum culture and may be associated with adverse treatment outcomes among tuberculosis patients. Accurately quantifying DC *Mtb* remains an important research objective, with current approaches tending to combine Most Probable Number (MPN) assays and culture filtrate (CF) supplementation. These assume that growth is equiprobable across all bacterial inoculum densities – an untested assumption for *Mtb*. We performed a half-logarithmic dilution series of *Mtb* from 70,000–0 CFU/mL, culturing each inoculum in either standard 7H9 or CF and monitoring growth by optical density. Inocula of ≥2,000 CFU/mL were 33 times more likely to grow than inocula of ≤700 CFU/mL. CF increased the odds of growth five-fold, and reduced the time-to-positivity by 294 hours (∼12 days) compared to 7H9 alone. However, CF’s growth-promoting effects diminished with increasing inoculum density, becoming negligible at 70,000 CFU/mL. Notably, CF broadly altered *Mtb* cell physiology, producing shorter bacilli that were less likely to incorporate the mycomembrane probe, DMN-trehalose. These data indicate that *Mtb* is poorly culturable from low inoculum densities – a limitation only partially overcome by CF. This non-uniform growth probability suggests that unsupplemented MPN assays may systematically underestimate *Mtb* CFU. Moreover, while CF promotes *Mtb* growth, its density-dependent activity and broader effects on cell physiology suggest that its influence extends beyond simply resuscitating DC *Mtb*. Improved methods are needed for detecting DC *Mtb* phenotypes, as these may support clinical care and shed light on factors that govern mycobacterial replication at different population densities.

**Highlights:**

- *Mycobacterium tuberculosis* (*Mtb*) culturability decreases at low inoculum densities
- Most Probable Number (MPN) assays may systematically underestimate *Mtb* CFUs
- Culture filtrate (CF) selectively promotes *Mtb* growth from low densities
- *Mtb* requires secreted factors in culture filtrate to initiate growth at low cell numbers
- CF-supplemented MPN assays may be inadequate for detecting DC *Mtb* phenotypes

## 1. Introduction

Tuberculosis (TB) remains a leading cause of death globally [1]. Culturing its infectious agent, *Mycobacterium tuberculosis* (*Mtb*), is essential to both laboratory research [2] and clinical diagnostics [3]. Even with molecular alternatives available [4], sputum culture continues to serve as the gold standard for diagnosing TB [3], monitoring treatment response [5], and performing drug susceptibility testing [6]. However, a notable limitation of *Mtb* culture is the prolonged turnaround time [6], driven by the bacterium’s inherently slow replication rate [7]. This limitation is compounded by the existence of differentially culturable (DC) *Mtb* phenotypes [8]: viable bacilli that retain their capacity to replicate, but grow poorly — or not at all — using standard solid or liquid growth media [9, 10]. DC *Mtb* have purported implications across the TB care cascade [11-13], including limiting diagnostic sensitivity [10, 14], reducing drug efficacy [15-17], and serving as hidden reservoirs for persistent infection [14, 18]. Their detection in sputum may also offer a biomarker for treatment monitoring [19]. As such, methods for the reliable detection and quantification of DC *Mtb* are critical.

Bacteria secrete several factors that promote growth, many of which act by resuscitating non-replicating phenotypes [8, 20]. In *Mtb*, such factors have been studied over many years, with Resuscitation-promoting factors (Rpfs) among the most well-characterised examples [21-23]. These murolytic enzymes degrade peptidoglycan to produce growth-promoting muropeptides [21, 22, 24, 25] and play an important role in both *in vitro* growth and virulence [23, 26, 27]. However, *Mtb* also possesses Rpf-independent mechanisms for growth resuscitation [9]. As a result, supplementing standard growth media with culture filtrate (CF) - sterile growth medium containing secreted cellular products including but not limited to Rpfs - may be sufficient for studying DC *Mtb* [18].

The simplest approach to detecting DC *Mtb* in clinical samples involves supplementing Mycobacterial Growth Indicator Tube (MGIT) media with CF and monitoring for changes in growth or time-to-positivity (TTP) [28, 29]. For quantification of DC *Mtb*, however, the Most Probable Number (MPN) assay is the preferred approach [8-10]. The MPN assay, widely used for microbial quantification in food and water safety, is a method that relies upon the Poisson distribution to estimate the number of culturable microorganisms in a sample by assessing growth across a series of dilutions [10]. When MPN assays are combined with CF supplementation, increased growth is often observed at the lowest inoculum densities relative to unsupplemented controls. These differences are attributed to the presence of DC *Mtb* [9, 10]. However, a critical assumption of this assay is that the probability of bacterial growth is uniform across all inoculum densities [30]. Here, we tested this assumption by systematically evaluating the culturability of *Mtb* from extremely low-density inocula. As detailed below, our results demonstrate that the probability of population growth is directly associated with initial cell density, therefore violating the key assumption. Instead, we show that CF partially restores growth at low densities, but its effect diminishes progressively with increasing bacterial burden.

## Methods

### 2.1 Strains and culture conditions

*Mycobacterium tuberculosis* H37Ra (ATCC 25177), referred to here as *Mtb*, was cultured in standard Difco™ Middlebrook 7H9 (BD Biosciences) supplemented with 10% BBL™ Middlebrook OADC Enrichment (BD Biosciences), 0.2% glycerol (v/v), and 0.05% Tween 80 at 37°C (7H9).

### 2.2 Stock titration

Titred stocks of *Mtb* were generated as previously described [31]. Briefly, *Mtb* cultures were prepared to an optical density at 600nm (OD_600_) of ∼ 0.1. Culture was diluted 1:1 in PBS + 0.1% Tween 80 and needle emulsified with sterile micro-emulsifying needles (CAD7974 Sigma Aldrich). Aliquots of the emulsified culture were then stained with SYBR Gold (ThermoFisher, S-11494) and Calcein-AM (ThermoFisher, C3100MP), respectively, to distinguish between metabolically active and permeable cells. In parallel, an aliquot was also heat-treated at 60°C for 12 minutes and stained with SYBR Gold to obtain a total cell count. Staining of media alone served as controls. All staining was performed for 60 minutes in the dark. Cell counts were acquired using the BD Accuri™ C6 flow cytometer and data analysis was conducted with the BD Accuri C6 software. Once the viable colony forming units (CFU) were enumerated, stocks were stored at -80°C.

### 2.3 Culture filtrate preparation

CF was prepared from *Mtb* cultures grown to an OD_600_ ∼0.4 and double filtered using 0.2 µm syringe filters (Merch Milipore, SLMP025SS). Fresh culture filtrate was prepared for each repeat.

### 2.4 Assessment of growth kinetics

Experiments were set up in U-bottom 96 well microtiter plates (Greiner Bio-One, 650 180). For each biological replicate, a frozen titred stock was thawed and diluted to a density of 1,000,000 CFU/mL, following which, a half-logarithmic (∼3.16-fold) dilution series was performed in sterile PBS; yielding approximately 300,000, 100,000, 30,000, 10,000, 3000, 1000, 300, 100 and 30 CFU/mL. Of each of the above dilutions, 10 µL were added to either 140 µL fresh standard 7H9 or fresh CF. The absolute CFU counts per well were 10,000, 3,000, 1,000, 300, 100, 30, ten, three, one and zero, with corresponding densities of 70,000 CFU/mL, 20,000 CFU/mL, 7,000 CFU/mL, 2,000 CFU/mL, 700 CFU/mL, 200 CFU/mL, 70 CFU/mL, 20 CFU/mL, 7 CFU/mL, and 0 CFU/mL. Given the accuracy of the initial stock titration and the relative nature of the growth comparison — growth in CF *vs*. standard 7H9 — these values were assumed to be accurate and were considered sufficient to address the research question.

The plate was incubated in a standing incubator at 37°C, and OD_600_ readings were acquired every 24 h (except over weekends) on a spectrophotometer (FLUOstar OPTIMA, BMG Labtech). These data were extracted and blank-normalised against the mean blank readings at each time point. Blank-corrected growth curves were smoothed with the loess function in R (span = 0.6) [32].

### 2.5 DMN-trehalose staining and microscopy

A starting *Mtb* inoculum of 20,000 CFU/mL was grown in either 7H9 or CF for approximately 11 days (to an OD_600_ of ∼0.5). The culture was split between two Eppendorf tubes, one of which was stained with 100 µM DMN-trehalose (DMN-tre) for two hours [33] while the other served as an unstained control. Samples were washed and resuspended in PBS for imaging on a Zeiss Axio Observer 7 equipped with a 100× plan-apochromatic phase 3 oil immersion objective (NA = 1.4). Epifluorescent illumination was provided by a 475 nm LED, and a Zeiss 38 HE filter set was used. Images were acquired using the Zeiss Zen software. Image segmentation and data extraction were done using MicrobeJ [34].

### 2.6 Measurements

#### 2.6.1 Growth outcomes

For each growth medium, the background signal (tolerance) was established from the range of OD_600_ values in uninoculated blank wells. For experimental wells, the mean and standard deviation of the first five OD_600_ readings (OD_start_) were calculated. A well was classified as positive for growth if the maximum OD_600_ reading exceeded OD_start_ by more than three standard deviations and the change in OD_600_ exceeded the blank tolerance range.

#### 2.6.2 Time-to-positivity

Time-to-positivity (TTP) was calculated from smoothed OD_600_ readings and defined as the time at which the maximum fold-change between two consecutive OD_600_ readings was observed. Fold-change was computed only when the OD_600_ exceeded three standard deviations above the blank and the OD_600_ gradient between two readings was >0.00. This prevented calculations when changes in OD_600_ readings were below the limit of detection. Although we originally intended to measure the time to maximal growth rate, limitations of OD-based detection at low densities required us to interpret TTP as the time until measurable exponential growth occurred.

#### 2.6.3 Doubling time

Doubling time was calculated using smoothed OD_600_ values and the following standard exponential growth equation:

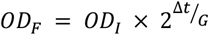

where OD_F_ and OD_I_ refer to the final and initial OD_600_ readings, respectively, and Δt and G refer to the time interval and doubling time, respectively. Rearranged:

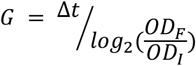

Owing to the challenges of detecting early growth in low-density cultures, the doubling time was estimated by averaging the three lowest values of G from each well, rather than by fitting a linear regression to log-transformed OD_600_ data.

#### 2.6.4 DMN-tre staining

Owing to the polar distribution of DMN-tre staining [33], maximum fluorescence intensity was used to determine stain uptake. A cell was classified as stained if its maximum intensity exceeded the average maximum intensity of the unstained controls by more than two standard deviations.

### 2.7 Data analysis

Binary outcomes (growth *vs*. no growth and DMN-tre staining) were assessed using logistic regression. For growth outcomes, a total of eight wells were assessed per inoculum density, per growth medium. Owing to the small sample size (n = 8; 2–3 per replicate), mixed effects logistic regression was not feasible. As such, a standard logistic regression model was fitted to evaluate the association between inoculum density (≥2,000 CFU/mL *vs*. ≤700 CFU/mL) and growth, adjusting for growth medium and assuming no major batch effects were present. For DMN-tre staining, standard logistic regression was used on representative data from one of two experimental repeats.

Linear regression was used to assess continuous outcomes (TTP, doubling time, DMN-tre intensity, and cell length). Mixed effects models were used to assess TTP and doubling time using the lme4 package [35] in R version 4.3.3 [36]. TTP was regressed against log_10_-transformed inoculum CFU and growth medium, while assessing the interaction between these two variables. Doubling time was regressed against a categorical CFU/mL (≥2,000 CFU/mL *vs*. ≤700 CFU/mL) and growth medium. In both models, experimental replicate was included as the random effect to determine effect sizes while accounting for batch effects. To investigate the association between DMN-tre intensity/cell length and growth medium, two separate linear regression models were run on representative data from one of two experimental repeats.

## 2. Results

### 3.1 Culture filtrate induced growth stimulation

To investigate the effect of CF on the culturability of *Mtb* from extremely low-density inocula, we performed a half-logarithmic (∼3.16-fold) dilution series of *Mtb* H37Ra from 70,000–0 CFU/mL (Figure 1A). Each inoculum was cultured in either standard 7H9 or CF and growth was monitored by optical density measurements using a microplate spectrophotometer. Growth outcomes (growth/no-growth), TTP, and minimum doubling time were calculated for each sample. The probability of sample contamination from external sources or from residual organisms in the CF was assessed by including at least 18 uninoculated blank wells per 96-well plate, for both 7H9 and CF. Across three replicates, this yielded a total of 64 blank wells per medium, none of which resulted in growth (Figure 1B).

**Figure 1:**
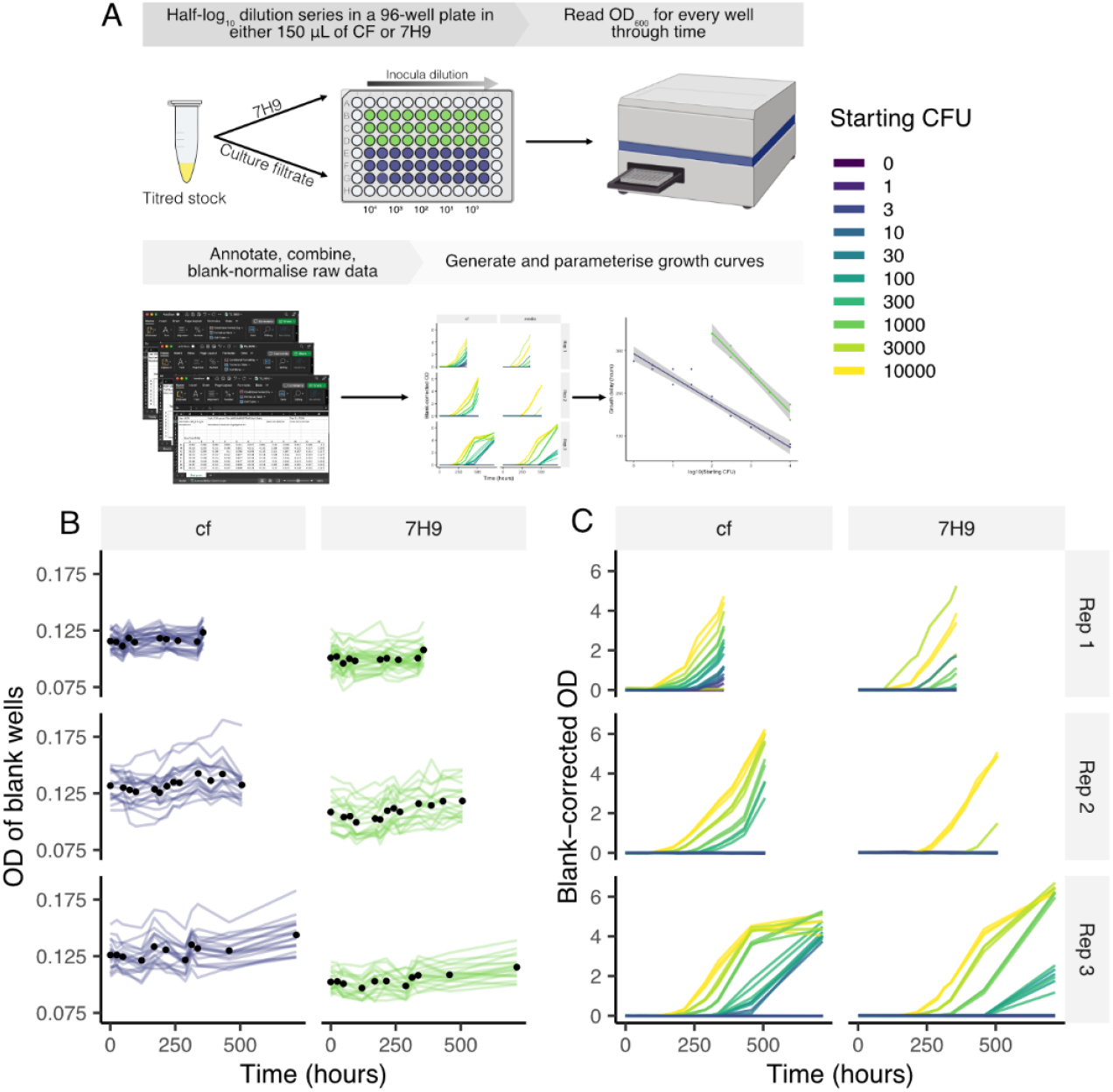
Culture filtrate induced growth stimulation. (**A**) Schematic representation of the experimental setup. Cells from an accurately titred freezer stock of Mtb were subjected to a half-logarithmic (3.16-fold) dilution series in either 7H9 (green) or culture filtrate (blue). The dilutions were added in triplicate (2/3 repeats) or duplicate (1/3 repeat) from column 2–11 on a 96-well microtiter plate with a final volume of 150 µL. All uninoculated wells were included as blanks. OD_600_ readings were recorded on a microplate spectrophotometer. Smoothed and raw blank-corrected OD_600_ readings were parametrised for further analysis. (**B**) OD_600_ readings from all blank wells through time across the replicates. The black circles represent the average blank reading per time point. (**C**) Representative blank corrected OD_600_ readings of Mtb grown from low inoculum densities (0–70,000 CFU/mL) in 7H9 OADC (7H9) or culture filtrate (CF). Each line represents one technical replicate (1/2) from one biological replicate. Rep = repeat. OD = OD_600_.

### 3.2 Culture filtrate partially improved poor culturability from low-density inocula

To generate a range of inoculum densities, we used *Mtb* freezer stocks that had been accurately enumerated by flow cytometry, ensuring precise and reproducible quantification of bacilli [31]. *Mtb* was poorly culturable from extremely dilute inocula, with a strong relationship observed between inoculum density and probability of growth (Figure 1C). While variability in growth was observed between biological repeats — which were conducted from independent serial dilutions of separate titred stocks — this was anticipated given the amplified impact of technical error at very low CFU. In this experiment, a gain or loss of only a handful of bacilli during titration would result in a fold-change in the final CFU. This batch variability was accounted for in our analytical approach. In contrast, variability in technical repeats (*i.e*., within batches) was low. Only two wells were identified as outliers — 7H9 inoculated with 3,000 and 10 CFU in the first replicate (Figure 1C top panel) — and these were excluded from subsequent analyses.

The probability of growth decreased with decreasing inoculum density for both CF and standard 7H9 (Figure 2A). No growth was observed in wells diluted to an expected final CFU of zero, supporting the accuracy of our dilution series and confirming that the concentrations of the initial inocula were low, as intended. Cultures initiated with inocula of ≥2,000 CFU/mL were 33 times more likely to grow compared to those with ≤700 CFU/mL, accounting for differences in growth media (Table 1). On average, CF increased the odds of growth five-fold, after adjusting for inoculum density. Together, these findings suggest that the probability of *Mtb* growth is dependent on inoculum density and is partially restored by CF at low initial CFU.

**Figure 2:**
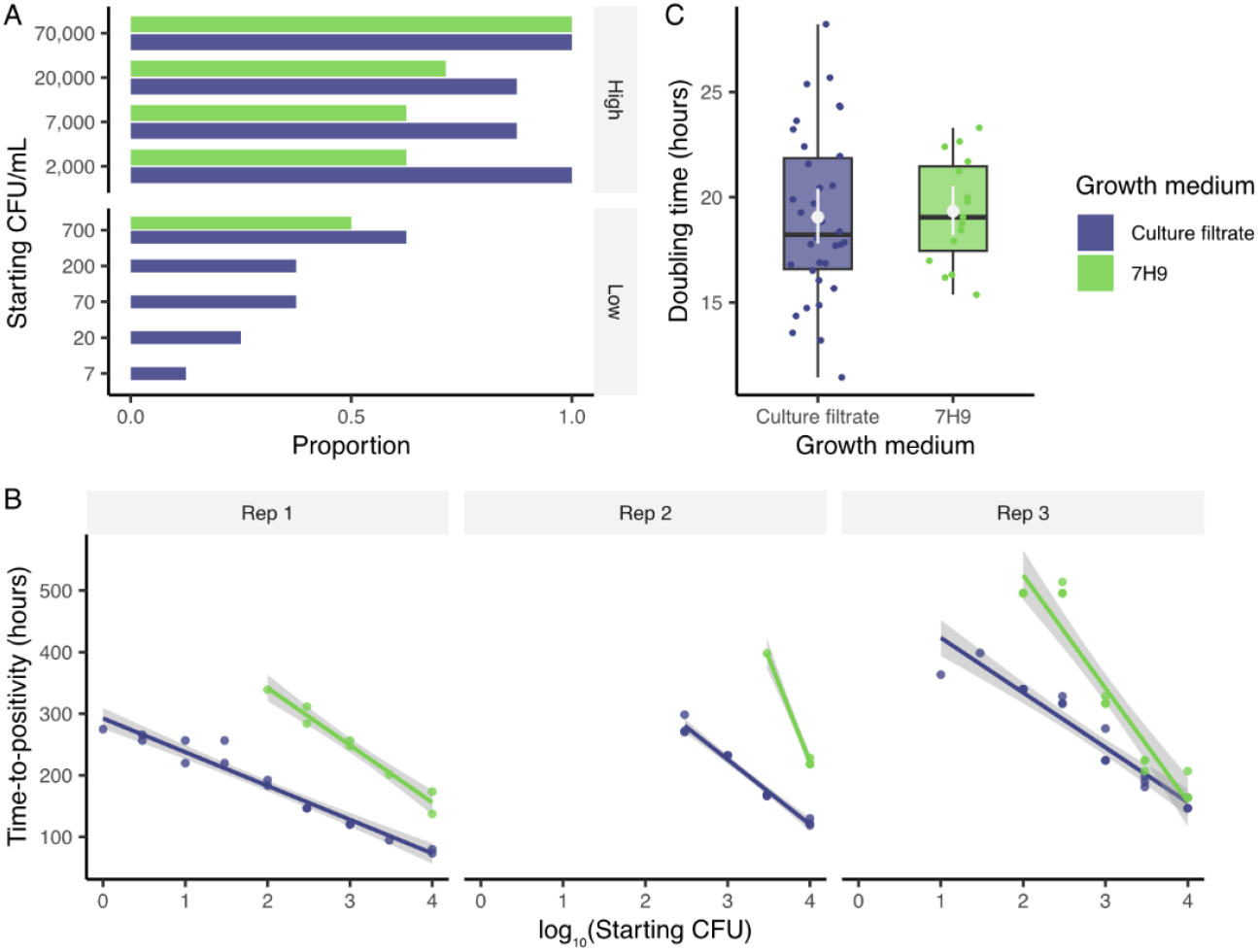
Culture filtrate partially improved poor culturability and reduced time-to-positivity from low-density inocula without impacting growth rate. (**A**) The proportion of wells demonstrating growth when inoculated in 7H9 (green) or CF (blue) stratified as high (≥2,000 CFU/mL) or low (≤700 CFU/mL) inoculum densities. Data are pooled from all three biological replicates, each with 2-3 technical replicates. (**B**) Time-to-positivity (in hours) plotted against the log_10_(absolute starting CFU), stratified by either 7H9 (green) or CF (blue) and by biological replicate (Rep 1–Rep 3). (**C**) The minimum doubling time for cells grown with or without culture filtrate. Data are pooled from all inoculum densities, as well as biological and technical replicates. The white circle and error bar represents the mean and 95% confidence interval, respectively.

**Table 1:** Culture filtrate partially improved poor culturability from low-density inocula. Results of a logistic regression comparing the odds of growth between starting CFU/mL (categorized as high [>700] and low [≤700]) and media (either or culture filtrate [CF]). Data are combined from three replicates.

|  | OR | P value | 95% CI |
| --- | --- | --- | --- |
| Starting CFU (ref. Low) | 33.1 | $>0.001$ | 13.1 ; 98.5 |
| Growth media (ref. 7H9) | 4.69 | 0.002 | 1.88 ; 13.4 |

### 3.3 Culture filtrate reduced time-to-positivity without impacting growth rate

*Mtb* growth was observed earlier for cells inoculated into CF compared to 7H9, with a more pronounced effect at lower inoculum densities (Figure 2C). We reasoned that this could reflect either a truncated lag phase or a faster doubling time. However, assessing the growth kinetics of *Mtb* from extremely paucibacillary inocula presented a challenge owing to the limited sensitivity of our optical density-based growth detection system, with no measurable changes in OD_600_ for several days (Figure 1C). Given this limitation, we were restricted to estimating the TTP — a commonly used correlate for inoculum density [37] — and the minimum measurable doubling time during exponential growth.

The TTP was linearly associated with the logarithm to the base ten of inoculum density (Figure 2C). On average, the TTP decreased by approximately 85 hours per log_10_ increase in starting inoculum, after accounting for differences between the growth media (Table 2). Culturing *Mtb* in CF further reduced the TTP by an average of 294 hours compared to 7H9 without reducing doubling time during exponential growth (Table 3). However, the reduction in TTP conferred by CF supplementation was not consistent across all inoculum densities (Figure 2C). Specifically, the difference in TTP between CF and 7H9 reduced by a further 58 hours for each log_10_ increase in inoculum density (Table 2), becoming almost negligible at inoculum densities above 70,000 CFU/mL (Figure 2C). These data suggest that CF supports *Mtb* growth by specifically reducing the lag phase of low-density inocula.

**Table 2:**
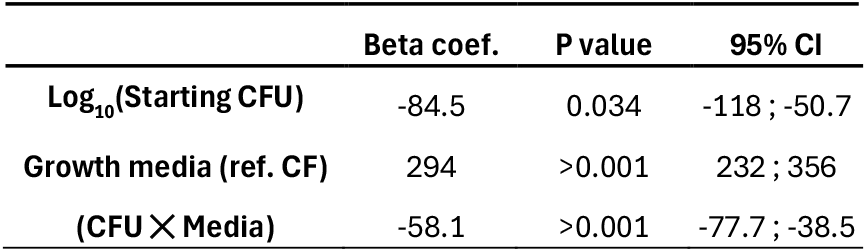
Culture filtrate reduced the time-to-positivity from low-density inocula. Results of a linear mixed effects model regressing time-to-positivity against log_10_(starting CFU), accounting for media (either 7H9 or culture filtrate [CF]), and the interaction between media and starting CFU. Data are combined from three replicates.

**Table 3:** Culture filtrate does not impact the doubling time during exponential growth. Results of a linear mixed effects model regressing doubling time against starting CFU/mL (categorized as high [>700] and low [≤700]) and media (either 7H9 or culture filtrate [CF]). Data are combined from three replicates.

|  | Beta coef. | P value | 95% CI |
| --- | --- | --- | --- |
| Starting CFU (ref. Low) | 1.06 | 0.414 | -1.46 ; 3.59 |
| Growth media (ref. 7H9) | 0.22 | 0.805 | -1.52 ; 1.97 |

### 3.4 Culture filtrate has long-lasting effects on cellular physiology

To investigate whether the effects of CF extend beyond growth initiation, we compared cell morphology and DMN-tre uptake between *Mtb* cultured with and without CF. Morphological variation in response to environmental stress is commonly observed in *Mtb* [38, 39], and DMN-tre is a fluorogenic probe incorporated into the mycobacterial outer envelope [40] with uptake influenced by growth phase [33] and stress [41]. Despite the importance of outer membrane remodelling during cell growth [42, 43], a proportion of *Mtb* bacilli remain unstained by DMN-tre during exponential growth, even after prolonged exposure to the fluorescent probe [40]. Notably, low-level DMN-tre uptake persists during stationary phase, suggesting ongoing maintenance of the outer envelope under conditions of reduced replication [33]. As such, we hypothesised that DMN-tre unstained cells may represent a non-growing phenotype.

Cultures grown in CF appeared less clumped than those in 7H9 — with an average ∼200 and ∼55 assessable single cells per image for CF and 7H9, respectively (Figure 3A & B, top panels). Surprisingly, DMN-tre incorporation was less efficient in CF (Figure 3C), staining 85.2% (2190/2571) and 98.5% (598/607) of cells grown in CF and 7H9, respectively (Table 4). Moreover, when stained, CF-grown cells exhibited markedly reduced fluorescence intensity (Figure 3D), being 0.26 log_10_(AFU) less bright compared to DMN-tre-stained *Mtb* grown in 7H9 (Table 5). Cell length was associated with both an increased staining efficiency (Table 4) and staining intensity (Table 5), and cells grown in CF were 0.10 log_10_(µm) shorter compared to those grown in 7H9 (Figure 3E, Table 6). Together, these findings suggest that CF has a broad and long-lasting effect on cell physiology that extends beyond growth initiation. While the functional implications of reduced DMN-tre uptake and shorter cell length remain unclear in the context of this work, these findings agree with previous observations [40] and strongly support the conclusion that CF broadly affects the whole population, influencing both growth initiation and downstream cellular physiology.

**Figure 3:**
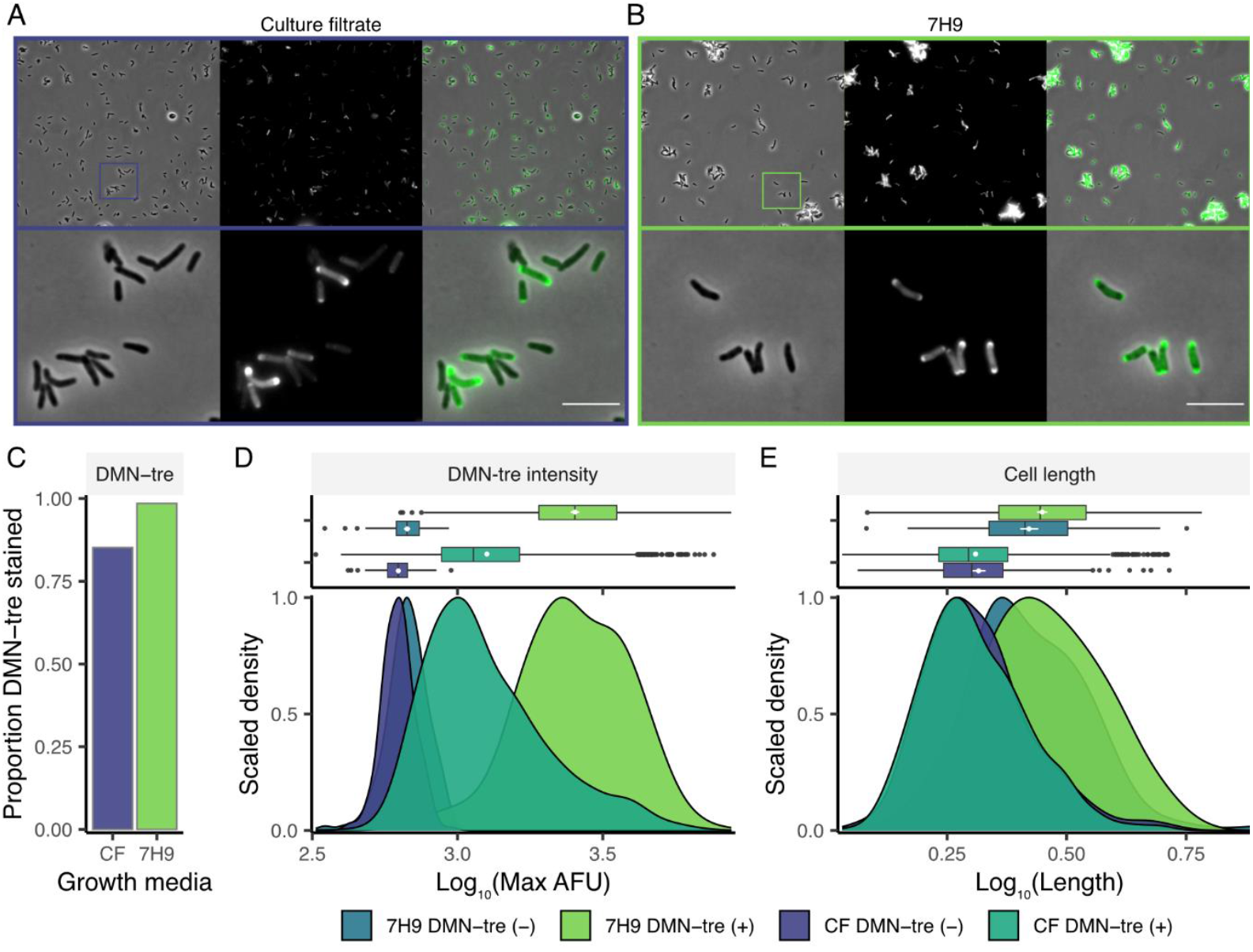
Culture filtrate has long-lasting effects on cellular physiology. Representative images of Mtb cultured in (**A**) culture filtrate (CF) or (**B**) 7H9 and stained with DMN-trehalose (-tre). From left to right, images are of phase contrast, DMN-tre fluorescence, and a merge of the two. The second row consists of images cropped and expanded from the first row. The scale bar is 5 µm. (**C**) The proportion of cells that stained with DMN-tre in CF (blue) and 7H9 (green). Box-and-whisker and equivalent density plots comparing the (**D**) maximum fluorescence intensity (in arbitrary fluorescence units [AFU]) and (**E**) cell length (log_10_[µm]) between cells cultured in 7H9 or CF and with (+) or without (-) DMN-tre. White circles and error bars on box-and-whisker plots represent the mean ±95% CI. Data are presented from one representative experiment (n = 2).

**Table 4:** Culture filtrate reduced odds of DMN-tre staining during exponential growth. Results of a logistic regression comparing the odds of staining with DMN-tre between growth media (either 7H9 or culture filtrate [CF]) after adjusting for cell length.

|  | OR | P value | 95% CI |
| --- | --- | --- | --- |
| Growth media (ref. 7H9) | 0.14 | >0.001 | 0.06 ; 0.25 |
| $\log_{10}(\text{Cell length})$ | 36.8 | >0.001 | 12.6 ; 111 |

**Table 5:** Culture filtrate reduced DMN-tre staining intensity during exponential growth. Results of a linear model regressing log_10_(maximum intensity) against growth media (either 7H9 or culture filtrate [CF]), adjusting for log_10_(length), DMN-tre staining, and the interaction between media and DMN-tre.

|  | Beta coef. | P value | 95% CI |
| --- | --- | --- | --- |
| Growth media (ref. 7H9) | -0.001 | 0.959 | -0.04 ; 0.04 |
| $\log_{10}(\text{Cell length})$ | 0.28 | >0.001 | 0.22 ; 0.34 |
| DMN-tre (ref. unstained) | 0.57 | >0.001 | 0.53 ; 0.61 |
| (Media × DMN-tre) | -0.26 | >0.001 | -0.31 ; -0.22 |

**Table 6:**
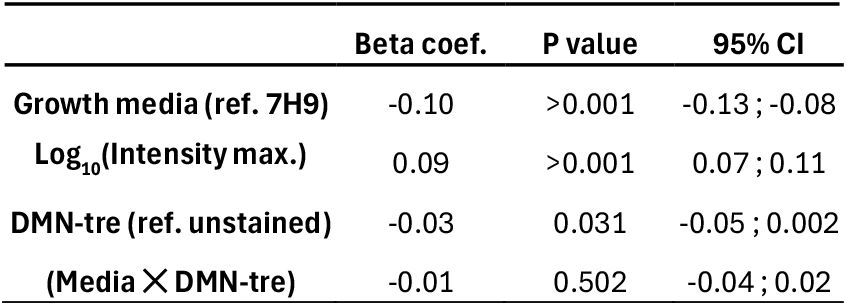
Culture filtrate reduced cell length during exponential growth. Results of a linear model regressing log_10_(cell length) against growth media (either 7H9 or culture filtrate [CF]), adjusting for log_10_(maximum intensity), DMN-tre staining, and the interaction between media and DMN-tre.

|  | Beta coef. | P value | 95% CI |
| --- | --- | --- | --- |
| **Growth media (ref. 7H9)** | -0.10 | >0.001 | -0.13 ; -0.08 |
| **Log<sub>10</sub>(Intensity max.)** | 0.09 | >0.001 | 0.07 ; 0.11 |
| **DMN-tre (ref. unstained)** | -0.03 | 0.031 | -0.05 ; 0.002 |
| **(Media × DMN-tre)** | -0.01 | 0.502 | -0.04 ; 0.02 |

## 3. Discussion

*Mtb* has long been associated with DC phenotypes [8]. These are proposed to have important implications across the TB care cascade [11] — including limiting diagnostic sensitivity [10], reducing drug efficacy [9, 15, 16], and serving as a potential biomarker for treatment monitoring [19]. These DC *Mtb* — later termed DCTB [10] — were first quantified in MPN assays through increased growth in wells supplemented with CF or CF derivatives, such as Rpfs, when compared to unsupplemented controls [8, 9]. While this is not the first study to highlight the flaws of this assay [44-46], here we sought to investigate a key assumption of this approach and its applicability to *Mtb*: namely, that the probability of bacterial growth is uniform across all dilutions such that no-growth wells are likely to be bacteria-free [30]. While this assumption may hold for other bacterial species, here, we show that it does not hold for *Mtb*, which becomes increasingly less likely to grow as inoculum density decreases. Furthermore, our data suggest that CF’s effect may not be limited to resuscitating DC phenotypes but may also enhance growth more broadly, particularly at low inoculum densities.

Quantifying *Mtb* CFUs at extremely low inoculum densities is challenging, complicating research involving paucibacillary samples. Regardless, our primary objective was to assess changes in growth between *Mtb* cultured in 7H9 *versus* CF. While the absence of growth at the lowest dilutions (<700 CFU/mL) might reflect a true lack of viable bacilli, the reproducible enhancement of growth in CF supports the conclusion that *Mtb* is poorly culturable from low-density inocula in standard media. Notably, CF’s growth-promoting effects were limited to the most dilute inocula, consistent with previous reports. For example, others have shown that low-density *Mtb* fails to grow in a microfluidic device unless exposed to CF [47] and, although tested at higher densities (20,000–60,000 CFU/mL), the growth-promoting effect of recombinant Rpf was strongest at the lowest densities [48]. Similarly, in MGIT-cultured clinical specimens, CF supplementation promoted growth from paucibacillary samples, including sputum from people living with HIV [28], patients on TB treatment [49], and cerebrospinal fluid from patients with spinal TB [29]. These findings suggest that *Mtb* may require secreted, bacterial-derived factors to initiate growth, which could explain CF’s diminishing utility as inoculum density increases without requiring the selective resuscitation of a distinct subpopulation. This density-dependent requirement for CF has important implications for the utility of MPN assays, which purportedly detect DC *Mtb* at the lowest dilutions when supplemented with CF. Although CF supplementation typically results in a predictable one to two log_10_ increase in estimated CFUs [9, 10] – an effect attributed to the presence of DC *Mtb* – our data suggest that this increase likely reflects CF’s partial restoration of growth at low inoculum densities. As such, the non-uniform, density-dependent effect of CF may introduce systematic bias, overestimating the number of DC organisms.

In addition to enhancing culturability from low-density inocula, growth in CF broadly altered *Mtb* physiology — changes that persisted into exponential growth. Compared to cultures grown in 7H9, CF-grown *Mtb* were shorter, appeared less clumped, and exhibited reduced uptake of DMN-tre — a marker of outer membrane metabolism [50]. While the mechanisms underlying these changes remain unclear, one possibility involves the role of Rpfs in peptidoglycan hydrolysis [21] and their synergy with other cell-envelope modifying enzymes [51, 52], which may indirectly affect outer membrane dynamics. This could account for the reduced DMN-tre uptake observed. Alternatively, while subtle differences in growth phase (*i.e*., being slightly earlier or later in exponential growth) could explain the morphological and staining differences observed here, we consider this to be unlikely. We selected a higher inoculum density (20,000 CFU/mL) for this experiment, where the effects of CF were less pronounced. Moreover, the observation that cells supplemented with CF stain less efficiently with DMN-tre is consistent with a previous report [40]. Additionally, growth in CF increases the concentration of Rpfs, which promote the activity of enzymes involved in cell division [52]. This could explain both the shorter cell length — comparable to previous observations from sputum [10] — and the reduced tendency to form clumps. Critically, CF induced changes in cell morphology and DMN-tre uptake throughout the population, supporting the conclusion that CF may not selectively resuscitate a non-culturable subpopulation but instead exerts broad physiological effects on *Mtb* bacilli.

Our study had several limitations that must be considered when interpreting the data presented here. A major limitation of this approach is the ecological fallacy: inferring individual-level parameters (*e.g*., growth of a subset of bacilli) from aggregate data (*e.g*., population expansion) [53]. This limitation is common to all studies using bulk measures of population expansion to understand the growth kinetics or (abundance) of individual cells, as is often the case in investigations of DC *Mtb* [9, 10, 14, 28]. While our primary study outcome was probability of growth at the population level, we can provide no additional insights into which phenotypes were — or were not — able to grow given the experimental conditions. While we used microscopy to partially overcome this limitation, time-lapse imaging is ultimately required to understand the growth outcomes of individual bacilli.

Reliably counting fewer than 10,000 *Mtb* CFU is technically challenging. Here, to best approximate the densities, a validated protocol based on flow cytometry was used to titre *Mtb* stocks [31]. Inaccuracies in stock titration and dilution might have impacted the exact densities used in this study and may explain the high variability between biological replicates. It should also be noted that freezing of *Mtb* results in a loss of culturability [9]. However, we don’t believe such deviations impacted the conclusions drawn here; any variability in exact counts or culturability was limited to biological replicates and was accounted for in our analytical approach. The complete absence of growth in wells with zero expected CFU, coupled with the reproducible linear relationship between TTP and estimated inoculum density, reinforces the reliability of our CFU estimates. Moreover, using OD_600_ to quantify *Mtb* growth from these extremely dilute inocula was suboptimal given its limited sensitivity; however, it facilitated the complexity of the experimental setup employed, enabling the comparison of several inoculum densities in different growth media. Consequently, this limitation precluded the accurate measurement of lag-phase duration and doubling time during exponential growth. As a result, we focused on two metrics: TTP and the minimum measurable doubling time — which we assumed would approximate the true minimum doubling time owing to the low density of the culture when changes in OD_600_ were detectable. Given the comparable doubling times observed between cultures in 7H9 and CF, we interpret the CF-associated reduction in TTP as evidence that CF shortens the lag-phase of *Mtb* from low-density inocula without affecting subsequent growth rates. An additional challenge arose due to suboptimal data acquisition, were non-uniform gaps in readings coincided with early exponential phase (mainly due to weekends). These gaps impacted the smoothing algorithm and reduced the accuracy of the doubling time approximation. They also impacted the TTP measurement, albeit to a lesser extent. Regardless, this error did not systematically bias results towards CF or 7H9 and should, therefore, only reduce the accuracy of our measurement without reducing the validity of the finding. Lastly, our focus on an expansive dilution series reduced the sample size per inoculum density. This limited our analytical approach, specifically regarding estimating the probability of growth from each inoculum density. Consequently, an arbitrary cutoff was selected to differentiate high (≥2,000 CFU/mL) and low (≤700 CFU/mL) densities and we were unable to quantify possible batch effects. In future, fewer dilutions will be tested to increase the number of replicates per plate. Regardless, the effect of CF on promoting growth was evident and is unlikely a consequence of the analytical approach employed here.

## Conclusion

While it might have been assumed previously, our study demonstrates that *Mtb* bacilli (even laboratory-adapted strains) are poorly culturable from low-density inocula. This finding has important implications for the use of MPN assays to quantify *Mtb* CFUs, as these rely on the assumption that growth is equiprobable across all inoculum densities. Our results suggest that the absence of growth at low MPN dilutions may reflect *Mtb*’s inherently poor growth efficiency from low-density inocula, rather than a high abundance of DC phenotypes. Consequently, while unsupplemented MPN assays likely underestimate *Mtb* CFU counts, CF supplementation introduces bias by predominantly restoring growth in the most dilute wells – resulting in a systemically increased CFU estimate. Although the existence of DC *Mtb* is well established and not disputed by our findings, we recommend caution in interpreting increased growth in CF-supplemented MPN assays or MGIT culture as direct evidence of DC phenotypes, as this is an example of the ecological fallacy. CF may simply enhance growth at low bacterial densities rather than resuscitate a distinct subpopulation. Further work is needed to develop methodologies to accurately quantify DC *Mtb*, as this may have important implications for clinical care and offer insights into the factors that govern *Mtb* replication at low population densities.

## CRediT authorship contribution statement

**Ryan Dinkele:** Conceptualization, methodology, software, formal analysis, writing – original draft, writing – review & editing, visualisation **Ditshego Ralefeta:** Investigation, writing – review & editing **Atica Moosa:** Investigation, writing – review & editing **Digby Warner:** Conceptualisation, resources, funding acquisition **Sophia Gessner:** Conceptualization, methodology, investigation, writing – original draft, writing – review & editing, supervising, project administration.

## Abbreviations

CF: Culture filtrate
CFU: Colony forming units
CI: Confidence interval
DC: Differentially culturable
DCTB: Differentially culturable tubercle bacilli
HIV: Human Immunodeficiency Virus
MPN: Most probable number
*Mtb*: *Mycobacterium tuberculosis*
OR: Odds ratio
Rpf: Resuscitation promoting factor
TB: Tuberculosis
TTP: Time-to-positivity

## Acknowledgements

The authors would like to thank Melissa Chengalroyen for her invaluable feedback, insightful comments, and constructive critique of this work.

